# Massive expression of germ cell specific genes is a hallmark of cancer and a potential target for novel treatment development

**DOI:** 10.1101/195305

**Authors:** Jan Willem Bruggeman, Jan Koster, Sjoerd Repping, Geert Hamer

**Affiliations:** Center for Reproductive Medicine, Amsterdam Research Institute Reproduction and Development, Academic Medical Center, University of Amsterdam, Amsterdam, The Netherlands.; Department of Oncogenomics, Academic Medical Center, University of Amsterdam, Amsterdam, The Netherlands.

**Author notes:** These authors contributed equally to this work (Jan Willem Bruggeman, Jan Koster).

## Abstract

Cancer cells have been found to frequently express genes that are normally restricted to the testis, often referred to as cancer/testis (CT) antigens or genes^1, 2^. Because germ cell specific antigens are not recognized as “self” by the innate immune system^3^, CT-genes have previously been suggested as ideal candidate targets for cancer therapy^4^. The use of CT-genes in cancer therapy has thus far been unsuccessful, most likely because their identification has relied on gene expression in whole testis, including the testicular somatic cells, precluding the detection of true germ cell specific genes. By comparing the transcriptomes of micro-dissected germ cell subtypes, representing the main developmental stages of human spermatogenesis^5^, with the publicly accessible transcriptomes of 2.617 samples from 49 different healthy somatic tissues^6^ and 9.232 samples from 33 tumor types^7^, we here discover hundreds of true germ cell specific cancer expressed genes. Strikingly, we found these germ cell cancer genes (GC-genes) to be widely expressed in all analyzed tumors. Many GC-genes appeared to be involved in processes that are likely to actively promote tumor viability, proliferation and metastasis. Targeting these true GC-genes thus has the potential to inhibit tumor growth with infertility being the only possible side effect. Moreover, we identified a subset of GC-genes that are not expressed in spermatogonial stem cells. Targeting of this GC-gene subset is predicted to only lead to temporary infertility, as untargeted spermatogonial stem cells can recover spermatogenesis after treatment. Our GC-gene dataset enables improved understanding of tumor biology and provides multiple novel targets for cancer treatment.

---

Where discovery of most CT-genes depended on whole testis expression data, we here used a unique list of genes expressed in human male germ cells generated in our laboratory^5^ to identify true GC-genes. Using R2, a genomics analysis and visualization platform we developed recently^8^, we compared these genes to data from the Genotype-Tissue Expression (GTEx) project^6^ and The Cancer Genome Atlas (TCGA)^7^. This comparison yielded 756 putative novel GC-genes (**supplementary data 1**). In order to visualize how the 756 GC-genes vary by tumor type, we stratified their expression in 33 tumor types in a heat map, showing that hundreds of GC-genes are expressed in all tumor types (**figure 1**).

**Figure 1.**
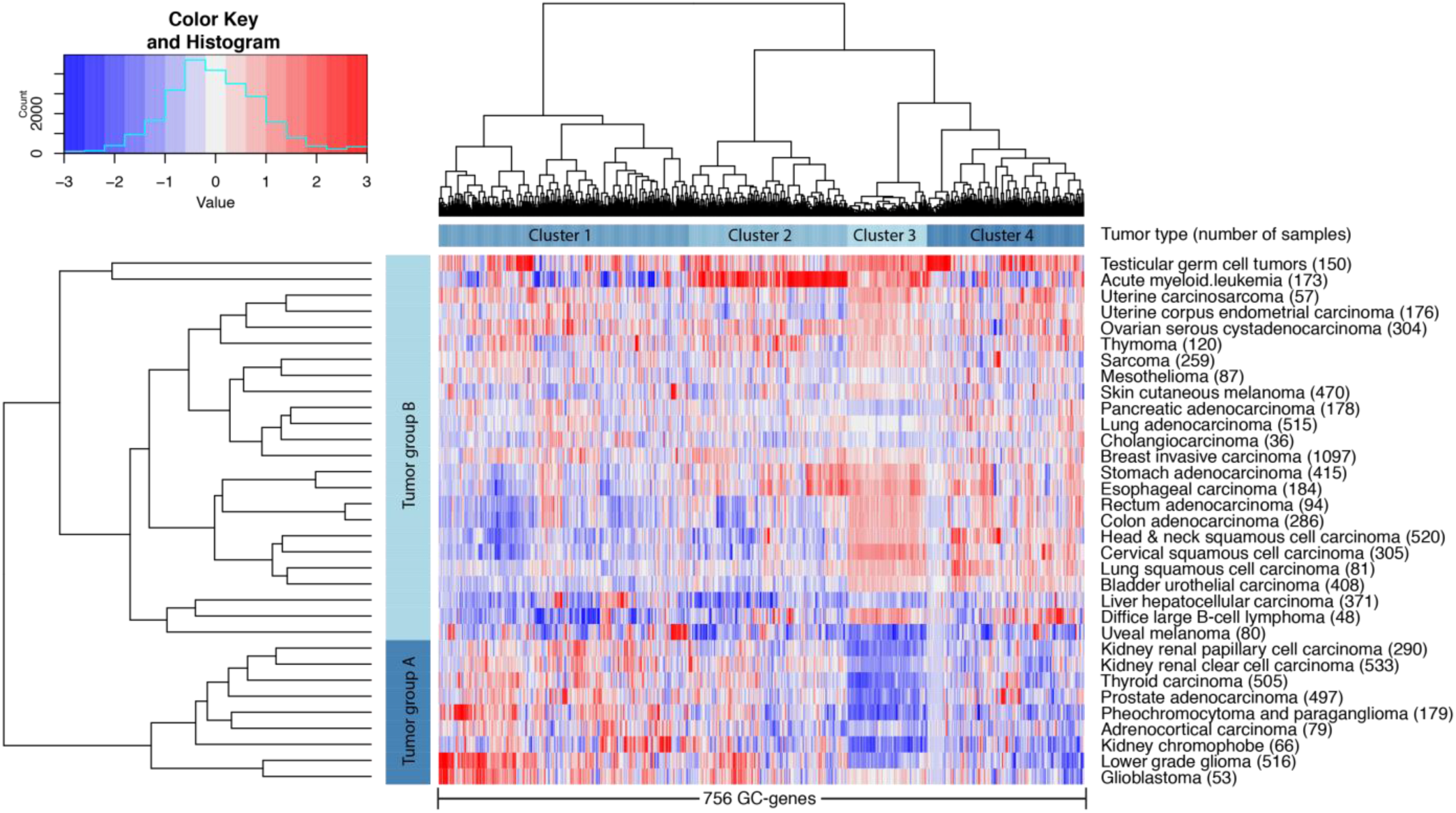
Hundreds of germ cell-specific genes are widely expressed in tumors. Shown here as hierarchical clustering of the average expression per tumor group (Euclidean distance, ward linkage). These germ cell cancer genes (GC-genes) divide tumors in two main groups, mainly based on GC-gene cluster 3, containing genes involved in mitotic and meiotic metaphase regulation. Gene expression levels are indicated by a Z-score dependent color, where blue and red represent low and high expression respectively.

For each of the three datasets, the maximum expression measured per gene was used to determine arbitrary inclusion criteria (**extended data 1**). To avoid false positive results, the selection criteria we applied to identify GC-genes are more stringent than previous selection criteria used to identify CT-genes. Moreover, whereas most studies allowed expression in 1-2 tissues other than the testis ^9-12^, our selection excludes all genes expressed in healthy tissues other than the testis. Thus, for most of the 756 genes identified in this study we can be certain that they are true GC-genes. However, lowly expressed genes that are only shortly or temporarily expressed, are only expressed in rare cell types, or only expressed under certain conditions may have escaped our selection. In addition, because germ cell tumors can be expected to express many germ cell specific genes, we analyzed which genes would not have been included in our initial list after exclusion of testicular germ cell tumors, and identified 45 GC-genes that are predominately expressed in germ cell tumors (**supplementary data 2**). From our original list of 16.589 genes expressed in male germ cells, 166 genes are present in a database containing genes specifically expressed in cancer and whole testis tissue, the cancer-testis(CT)-database^13^. From the 255 CT-genes in this database, only 26 overlap with our newly identified 756 GC-genes. This can be explained by the fact that the testis mostly consists of somatic cells. Germ cell specific RNAs can therefore be diluted below detection levels in whole testis lysates, while testicular somatic genes are not included in our analysis. Indeed, from a more recent analysis that revealed 1019 potential CT-genes^14^, only 117 (13%) were also present in our analysis (**figure 2**). These data combined, our current analysis has identified 630 GC-genes that have not been previously identified as CT-genes, 615 of which are expressed in non-testicular tumors.

**Figure 2.**
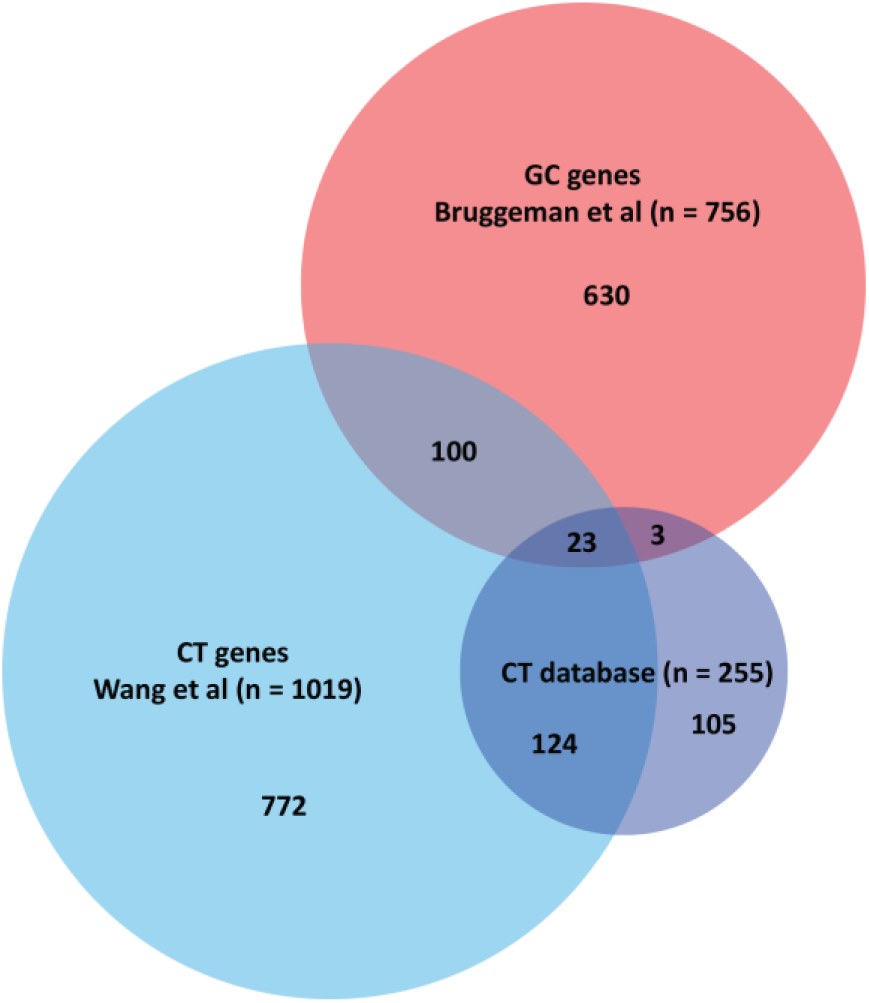
Most GC-genes have not been described before as CT-gene. Venn diagram comparing the present analysis of germ cell-specific cancer (GC) genes (red) to earlier identified Cancer/Testis (CT) genes by Wang et al (light blue) and the CT-database (dark blue). The number in each section represents the number of genes.

Hierarchical cluster analysis revealed that, based on expression of GC-genes, the tumors form two main groups, mostly characterized by high or low expression of a specific subset of GC-genes (gene cluster 3) (**figure 1 & supplementary data 3A**). Interestingly, gene ontology (GO) analysis using DAVID Bioinformatics Resources v6.7^15^ revealed that this gene cluster predominately contains genes involved in M-phase and cell cycle regulation, intriguingly both mitotic and meiotic (**table 1**). Also processes pivotal to meiosis, such as DNA double-strand break repair and homologous recombination, are well represented in this cluster (**supplementary data 3D**). Further GO-analysis revealed that six biological processes are significantly represented by all 756 GC-genes (**supplementary data 3F**): the regulation of transcription and gene expression, including the metabolic processes required for RNA and DNA synthesis, the M-phase of the mitotic and meiotic cell cycle, DNA double-stranded break repair, DNA metabolic processes, spermatogenesis and cell adhesion. Additional gene ontology analysis was performed on the top 25% GC-genes that were most widely expressed in tumors. Six biological processes appeared to be significantly represented by these 189 GC-genes, including cell cycle regulation and checkpoints, post-translational protein modification and DNA damage responses (**supplementary data 3G**). In line with previous research^14,16,17^, these processes suggests that GC-genes are not just randomly expressed germ cell specific genes but may actually contribute to tumor cell survival, proliferation and metastasis.

**Table 1.**
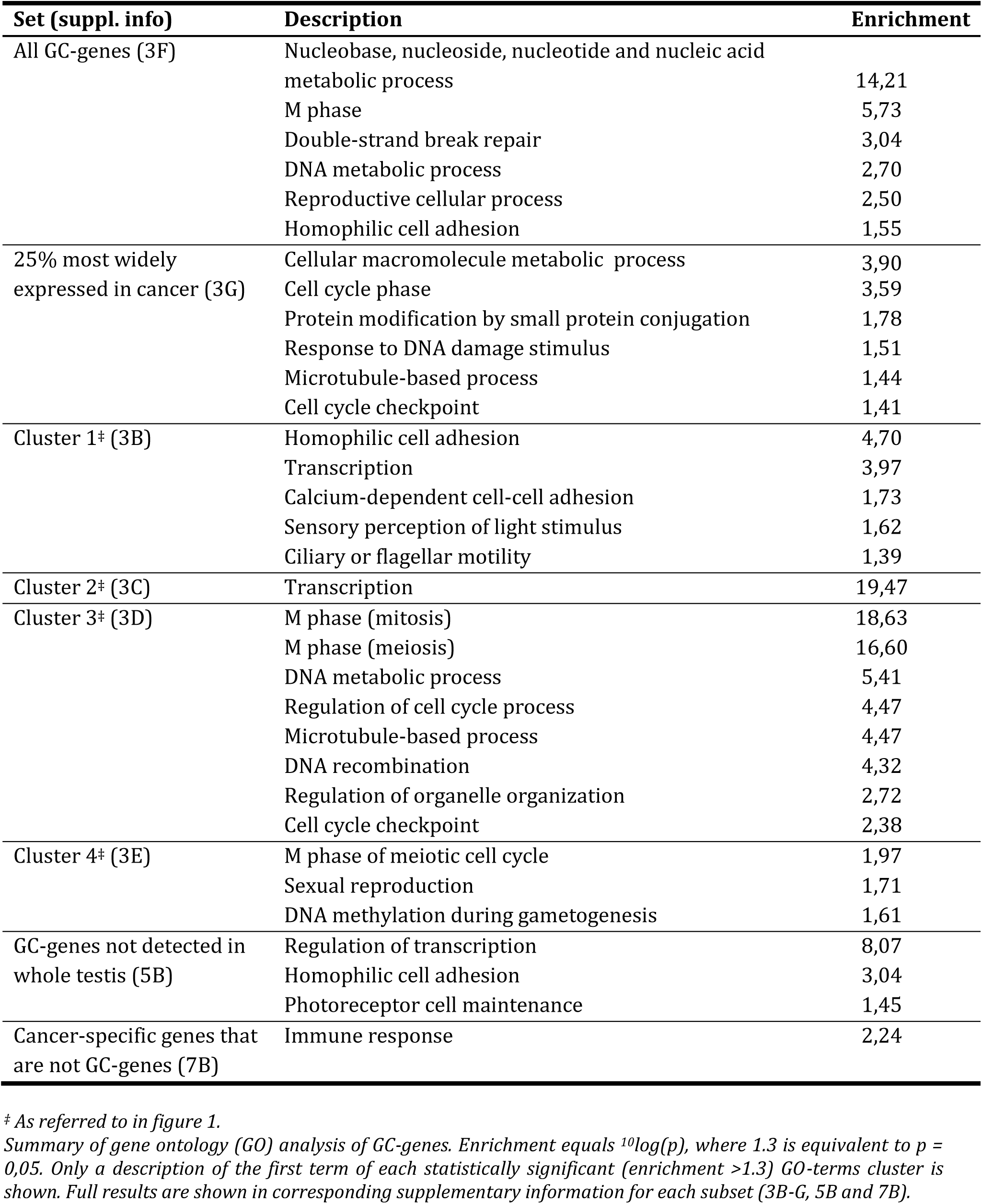
GC-genes represent processes that are likely to contribute to tumor cell survival, proliferation and metastasis.

Because proteins that are located on the outer cell surface would be ideal targets for induced adaptive immune (therapeutic) responses, we used the Panther 10.0 classification system^18^ to identify 113 GC-genes that encode plasma membrane proteins (**supplementary data 4A**). Seven proteins (GGTLC2, GP1BA, IGLL1, IL12RB2, NLGN1, NRG1 and UMODL1) are predicted to be located on the external side of the plasma membrane, of which two (BP1BA and GGTLC2) are anchored to the plasma membrane. Although highly expressed in some tumors, these seven genes are not expressed in most tumor types. The remaining 105 proteins are not predicted to be located on either side of the plasma membrane, and may or may not be suitable therapeutic targets. For 26 membrane protein-encoding genes, the average expression is high in all 33 analyzed tumor types (**supplementary data 4B**).

Because CT-genes have previously been identified using gene expression profiles of whole testis, including the testicular somatic cells, we additionally compared gene expression of whole testis tissue from the GTEx project^6^ to gene expression in human male germ cells^5^. From all genes with significant expression in male germ cells and no significant expression in any other tissue^6^, the difference in expression in comparison to whole testis was calculated. In order to correct for using different expression distributions, genes with difference below one^2^log value were excluded. This resulted in a list of 706 genes that are expressed in germ cells, but were not previously detected in testis as a whole. When comparing our list of 756 GC-genes to these 706 genes, we identified a subset of 334 GC-genes (44%) whose expression is very low or undetectable in testis as a whole (**supplementary data 5A**). Interestingly, among this subset of 334 GC-genes, the average expression in tumors is higher than in the remaining GC-genes (p=0.029, two-tailed T test). GO-analysis of this subset identified transcription regulation and cell adhesion as significantly enriched processes (**supplementary data 5B**), both essential for germ cell development, tumor proliferation and metastasis. This, and the fact that they are highly germ cell and cancer specific, makes these genes interesting candidates for future research.

Previously, CT-genes have been divided in X and non-X CT-genes, depending on whether they are located on the X chromosome or not. According to the CT-database, approximately half the CT-genes are CT-X^13^. However, of the 1019 CT-genes identified by Wang et al., only 105 were located on the X chromosome^14^. In line with this study, our analysis returned only 29 (4%) X-linked GC-genes and GC-genes seem to be distributed evenly across all chromosomes (**extended data 2**).

To validate to what extent germ cell specific RNA expression reflects protein expression in various human tissues we used the Human Protein Atlas (v15)^19^. From the atlas we retrieved all proteins expressed in testis or ovary and selected for highly reliable immunohistochemistry (Premium Tissue). In addition, because many CT-genes are known to be expressed in trophoblasts^20^ that will later form the placenta, we similarly retrieved all proteins expressed in placenta. The resulting genes (proteins) were then aligned with our 756 GC-genes, resulting in a list of 49 genes that were manually checked for germ cell– or placenta specific protein expression (**supplementary data 6**). This yielded three proteins that are exclusively present in placenta and 24 proteins that are present in male germ cells (and not in somatic cells of the testis or elsewhere). Of these, PRSS21 may be of particular interest, as it appeared to be a putative outer cell membrane protein and thus a possible therapeutic target.

To develop a therapy without side-effects in healthy tissues it would in principle be sufficient to identify genes that are uniquely expressed in tumors. For this, our list of human germ cell expressed genes would not be required. We therefore performed a similar analysis without our list of germ cell expressed genes and including testis from the GTEx database. This resulted in 724 cancer-specific genes, of which 301 genes appeared not to be GC-genes (**supplementary data 7A**). GO-analysis revealed that these 301 genes are predominately involved in immunological responses (**supplementary data 7B**). Hence, in contrast to germ cell specific genes, targeting these genes as a cancer therapy can be expected to lead to immunological side-effects.

Infertility is a major side effect of current anticancer treatments and would still be a potential side effect when targeting most GC-genes. A way to circumvent this would be to exclude genes expressed in the spermatogonial stem cells. In humans, these stem cells are included in the pool of quiescent or mitotically proliferating and differentiating spermatogonia, and are required to maintain life-long spermatogenesis. Because our dataset contains information about germ cell type-specific gene expression^5^, we were able to exclude genes expressed in spermatogonia. Of the 756 GC-genes, 69 displayed negligible expression in the spermatogonial stages (**supplementary data 8**). Hence, targeting these 69 GC-genes would not affect the spermatogonial stem cells and therefore only lead to temporary infertility. Importantly, we have recently found that spermatogonia already express many mRNAs that are not translated until later stages during spermatogenesis^5^. This implies that the number of GC-genes that can be targeted without inducing permanent infertility will most likely be larger than 69.

We here show that expression of hundreds of germ cell specific genes may not only contribute to already established hallmarks of cancer^21^, but can be considered as a hallmark of cancer in itself. Germ cells and cancer cells share the intrinsic drive to propagate, regardless of survival of the soma^22^-^24^. Studying the behavior and characteristics of germ cells may thus lead to novel insights in cancer development. Because our datasets are publically available, more tumor types can now be analyzed on the expression of germ cell specific genes. We anticipate that this will lead to a better understanding of tumor biology and improved treatment options.

## Methods

In order to identify genes whose expression is restricted to germ cells, we compared a list of genes expressed in human male germ cells generated in our laboratory^5^ with a publicly accessible dataset from the Genotype-Tissue Expression (GTEx)^6^ project containing the transcriptome of 2.921 samples of 53 healthy non-cancerous tissue types. From this dataset we excluded ovary, testis and two transformed cell lines. This yielded 2.617 samples from 49 different tissue types (**supplementary data 9A**). For gene expression in cancer we used a publicly accessible dataset from The Cancer Genome Atlas (TCGA)^7^, containing the transcriptomes of 9.232 samples from 33 tumor types (**supplementary data 9B**). To identify genes that are exclusively expressed in germ cells and at least one tumor type, these three sources were combined using R2, a genomics analysis and visualization platform we developed recently^8^ (**supplementary data 1**).

From the 16.589 adult male germ cell genes, 1.526 genes were excluded for which no information was available on the expression in either non-cancerous somatic tissues (GTEx)^6^ or tumors (TCGA)^7^ (**supplementary data 10**). For gene expression in germ cells, genes with a maximum expression below 1.6 on a^2^log scale were considered background noise and were excluded (**extended data 1A**). Likewise, in order to only include genes that are exclusive to the male germ cells, genes with an expression over 1.8 on a^2^log scale in any non-cancerous somatic tissue were also excluded (**extended data 1B**). Finally, we selected for genes with an expression higher than 6.2 on a^2^log scale in at least one of 33 tumor types (**extended data 1C**).

The overlap between the CT database, Wang et al. and the present analysis was assessed by converting gene names to one common annotation (**supplementary data 11A-C**). 21 out of 276 genes in the CT database were either merged with existing genes (n=19) or could not be retrieved (n=2) (**supplementary data 11D**). Figure 2 was created using Biovenn (Hulsen, T., et al. 2008, *BMC Genomics* **9**: 488).

## Supplementary data

Supplementary data are available on request.

## Acknowledgements

This work was supported by ZonMw VIDI-grant 91796362 to S.R., an AMC Fellowship and The People Programme (Marie Curie Actions) of the European Union’s Seventh Framework Programme (CIG 293765) to G.H..

## Author Contributions

J.W.B, J.K. and G.H. conceived and designed the study. J.W.B, J.K. and G.H. performed bioinformatic analyses. J.W.B. and J.K. and performed data visualization. J.W.B., J.K. and G.H. interpreted the results. J.W.B, J.K., S.R. and G.H. critically read and wrote the manuscript.

